# α-Melanocyte Stimulating Hormone Reduces Blood Glucose Across Species

**DOI:** 10.1101/2025.03.26.645414

**Authors:** Patrick Swan, Brett Johnson, Stephanie E. Simonds, Tomris Mustafa, Michael J. Houghton, Suhaniya Samarasinghe, Neil G. Docherty, Migena Luli, Jack T. Pryor, Carissa Greenough, Anne Gibbon, Philip E Thompson, Wenxiao K Yue, Giuseppe de Vito, Gary Williamson, Carel W. le Roux, Alexander D. Miras, Michael A. Cowley

## Abstract

Peripheral administration of α-melanocyte stimulating hormone (α-MSH) promotes glucose uptake in skeletal muscle via a melanocortin 5 receptor (MC5R)-dependent mechanism in rodents. Here we demonstrate that this pathway is functional in humans. During an oral glucose tolerance test (GTT) in healthy human volunteers, peripheral administration of α-MSH was well tolerated, and reduced glucose and insulin incremental area under the curve by 39% and 35%, respectively. During a GTT in non-human primates, both α-MSH and a selective MC5R agonist, PG-901, increased glucose tolerance. *In vitro*, both α-MSH and PG-901 induced glucose uptake in primary human and non-human primate myotubes and immortalised human myotubes. Subcutaneous injection of PG-901 reduced blood glucose concentrations during a GTT in WT mice but had no effect in MC5R deficient mice. Thus, the novel MC5R-skeletal muscle pathway found in animals is functional in humans and represents a novel target for glycaemic control in people with diabetes.

## Main

The melanocortin system comprises five G protein-coupled melanocortin receptors (MCRs). Endogenous ligands of the MCRs include adrenocorticotropic hormone (ACTH), agouti-related protein (AgRP) and the melanocortin neuropeptides α-melanocyte stimulating hormone (α-MSH), β-MSH, γ-MSH^1^. The action of the central melanocortin system is well characterized, wherein MC3R and MC4R signaling regulates energy balance, growth, onset of puberty and food intake ^2–4^. In peripheral tissues, the function of the melanocortin system as it pertains to substrate metabolism is less well understood. The evidence for MCR signaling in skeletal muscle fatty acid oxidation, adipocyte lipolysis, insulin sensitivity, and peripheral glucose metabolism, to date only exists in animal models^5–12^.

We previously discovered that in animals, the peripheral actions of α-MSH, a non-selective agonist at melanocortin receptors 1, 3, 4, and 5, reduces plasma glucose via MC5R-mediated skeletal muscle glucose uptake^13,14^. MC5R, the only MCR expressed in rodent skeletal muscle, agonism by α-MSH does not trigger translocation of GLUT4 and thus this mechanism is independent of insulin-mediated glucose uptake^14^. We have also demonstrated that simultaneous activation of the MC5R pathway with α-MSH and the Akt-GLUT4 pathway with insulin have additive effects on glucose uptake in mouse vastus lateralis and extensor digitorum longus muscles ex vivo^14^. The functionality of melanocortin signaling in skeletal muscle, more specifically the MC5R-mediated glucose uptake pathway has not previously been explored in human biology.

Herein we use a series of interlinked experiments, including first-in-human experimental medicine studies, in vivo, and in vitro approaches to test two hypotheses: (i) the physiological effects of improved glucose tolerance in response to peripheral melanocortin receptor agonism is preserved across species, including humans, and (ii) the physiological mechanism underpinning this effect involves increased skeletal muscle glucose uptake downstream of MC5R agonism.

## Results

### Human studies

Two cohorts of healthy participants were recruited for a dose-finding and a replication study. All studies were, placebo controlled, double-blinded, and the order of infusions randomized. Infusions were initiated 30 minutes prior to the OGTT. The exploratory dose-finding study was designed to assess safety and tolerability and identify the dose with the maximal effect on plasma glucose lowering. In the dose-finding study, eight male and seven female participants (mean age of participants was 28.0 ± 8.4 years, BMI 25.0 ± 3.0 kg/m^2^ and HbA1c 32.0 ± 4.1 mmol/mol (**Supplementary Table 1**)). Participants were infused with low (15 ng/kg/hr), medium (150 ng/kg/hr), or high (1500 ng/kg/hr) doses of intravenous α-MSH and saline during OGTT on four different study visits. During hyperinsulinaemic clamp visits, participants were infused with the medium dose of α-MSH or saline in a placebo controlled, double-blinded, randomized manner.

### Oral Glucose Tolerance Tests in humans

In the dose-finding cohort, there was a dose-dependent lowering of plasma glucose concentration in response to increasing doses of α-MSH with the high dose corresponding to a 24.6% reduction in glucose iAUC_0-120_ compared to saline (**Supplementary Table 2)**. The corresponding reduction of plasma glucose concentration compared to saline with the low and medium doses of α-MSH was 6.0% and 12.9% respectively (**Supplementary Table 2)**. The high dose of α-MSH infusion reduced insulin iAUC_0-120_ by 18.4% compared to saline, with the low dose reducing serum insulin by 11.9% and the medium dose increasing it by 4.2% (**Supplementary Table 2)**.

### Hyperinsulinaemic-Euglycaemic Clamps in humans

In the dose-finding cohort a modest increase in glucose infusion rate (GIR) was observed during 150ng/kg/min α-MSH infusion compared to saline (**Supplementary Figure 2)** although this did not reach statistical significance.

### Oral Glucose Tolerance Tests-Replication Study

The replication study was powered based on the results of the dose-finding study and designed with the aim of confirming its findings in a different cohort of 22 healthy volunteers, eleven males and 11 females (mean age of participants 29.1 ± 9.1 years, BMI 22.7 ± 3.1 kg/m^2^ and HbA1c 33.0 ± 0.8 mmol/mol (**Supplementary Table 1))** were infused with α-MSH (1500 ng/kg/hr) or saline during an OGTT on two different study visits using a double-blinded, randomised approach.

In the replication study, oral glucose increased glucose levels in all participants, infusion of high dose α-MSH significantly reduced plasma glucose iAUC _0-120_ by 39.1% compared to saline (**Figure 1.A-B**). The reduction in plasma glucose concentrations during the OGTT was statistically significant 30 (7.6±0.29 mmol/L vs. 6.5±0.26 mmol/L) and 60 (7±0.4 mmol/L vs. 5.6±0.3 mmol/L) minutes post glucose bolus (Figure 1A,). A concomitant 35.7% reduction in serum insulin iAUC _0-120_ was observed during high dose α-MSH infusion compared to saline (**Figure 1.C-D**). The reduction in serum insulin concentrations during the OGTT was statistically significant at time points 60 and 120 minutes (Figure 1C).

**Figure 1.**
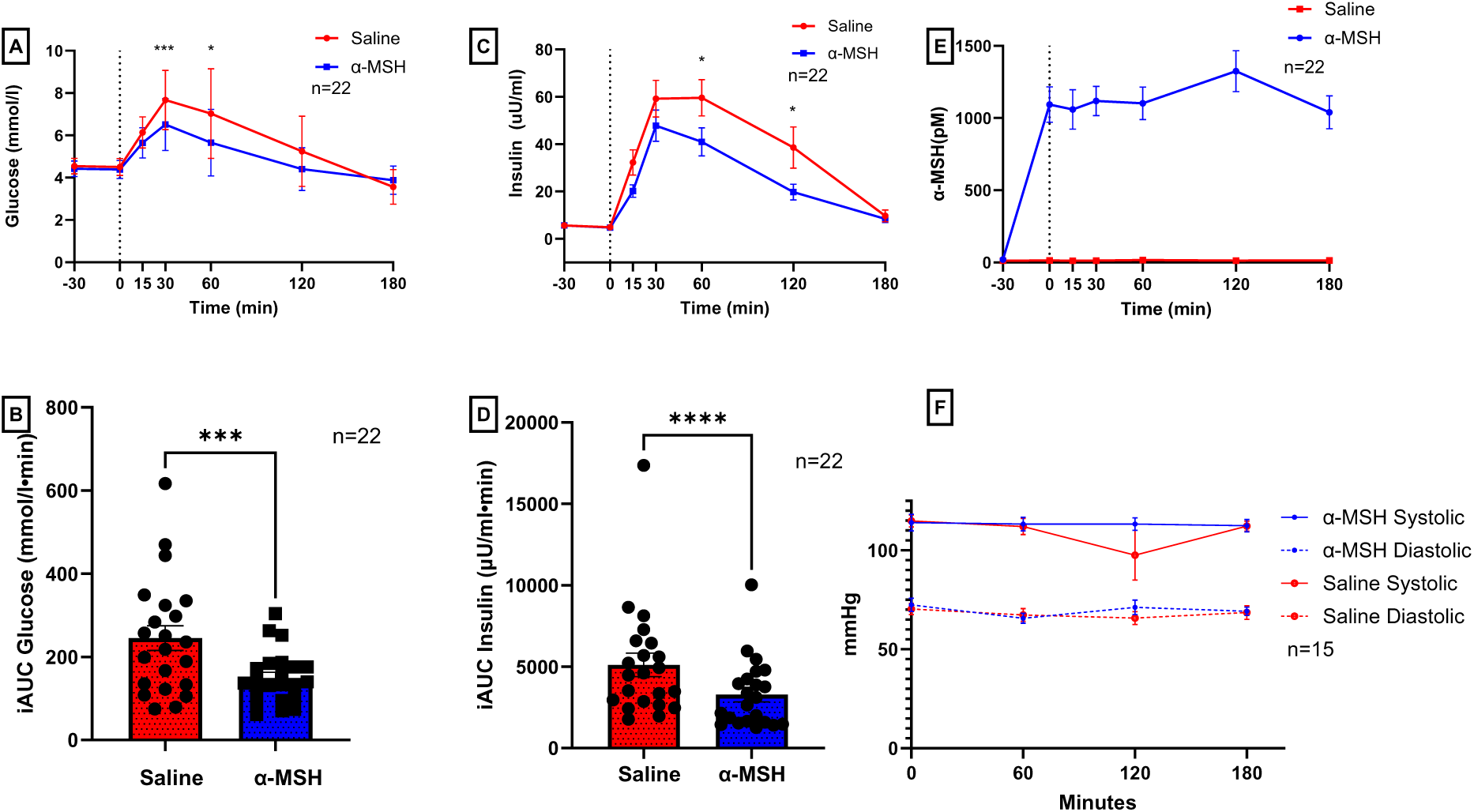
Effect of intravenous infusion of a-MSH in healthy humans. Data are presented as mean ± SEM. Timepoint of oral glucose administration marked by dotted line. **A-B)** 1500 ng/kg/hr α-MSH significantly reduced plasma glucose concentrations during the OGTT at 30 and 60 minutes, (replication cohort n=22, two-way ANOVA time x infusion *p=0.015), and reduced plasma glucose concentration iAUC_0-120_ by 39.1% compared to saline. (replication cohort n=22, paired t-test ***p<0.001). **C-D)** significantly decreased serum insulin concentrations at 15, 30 and 60 minutes during the OGTT (replication cohort n=22, two-way ANOVA time x infusion *p=0.01) and reduced serum insulin concentration iAUC_0-120_ by 35.7% (replication cohort n=22, paired t-test ****p<0.0001). **E)** Acute intravenous infusion of α-MSH (1500 ng/kg/hr) rapidly increased plasma α-MSH concentrations that remained stable during the duration of infusion (OGTTs, replication cohort n=22), **F)** 1500 ng/kg/hr α-MSH did not have a significant effect on systolic and diastolic blood pressure (dose-finding cohort, n=15).

In the replication cohort, circulating α-MSH concentrations remained stable throughout the OGTT at 15.8± 1.1 pM during saline infusion. α-MSH concentration elevated to 1092 ± 122.8 pM after 30 minutes of α-MSH infusion at 1500 ng/kg/hr, at which time the OGTT was performed. α-MSH concentrations remained stable (1039-1324 pM) for the duration of the OGTT (**Figure 1.E)**.

### Safety Profile in humans

At all doses tested infusion of α-MSH was well tolerated without evidence of nausea, flushing, or any other adverse events. There was no significant difference in the incidence of hypoglycaemia (plasma glucose ≤3.9 mmol/L) between saline and high dose α-MSH infusions (0.03% vs 0.06%, respectively). Low (15 ng/kg/hr) and medium (150 ng/kg/hr) doses of α-MSH demonstrated a similar incidence of hypoglycaemia across all timepoints (0.03% and 0.04%, respectively) (**Supplementary Table 3**). α-MSH did not have a significant effect on systolic and diastolic blood pressure, (**Figure 1. F)** or heart rate (**Supplementary Figure 3)** at any timepoint.

### Glucose tolerance tests in non-human primates

Male southern pigtail macaques (*Macaca nemestrina*) were anaesthetized and then treated with intravenous infusions of saline or α-MSH (0.167 or 0.334 mg/kg/hr) followed 30 min later by an intravenous glucose tolerance test (600mg/kg). Glucose infusion increased glucose levels in all monkeys, α-MSH infusion resulted in a non-significant dose-dependent increase in glucose tolerance, with a 29% decrease in the iAUC of glucose between saline and 0.334 mg/kg/hr dose of α-MSH (Figure 2. A-B).

**Figure 2.**
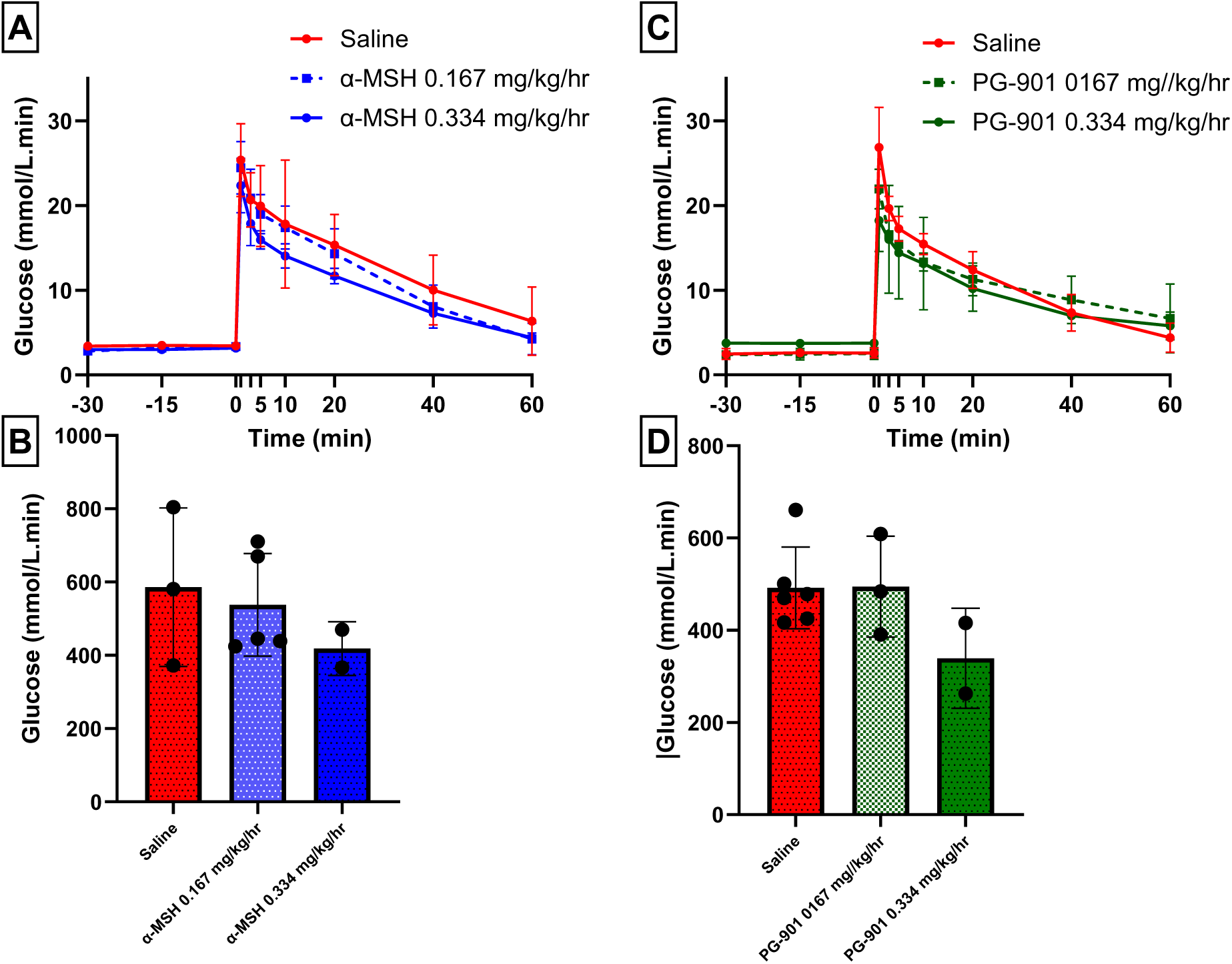
Effect of intravenous infusion of α-MSH and PG-901 in non-human primates. Data are presented as mean ± SD. **A-B)** In male macaques, intravenous infusion of α-MSH at 0.167 mg/kg/hr (n=5) and 0.334 mg/kg/hr (n=2) reduced plasma glucose concentrations and glucose iAUC during an IV glucose tolerance test compared to saline (n=3) **C-D)** In female macaques, intravenous infusion of PG-901 at 0.334 mg/kg/hr (n=2) reduced plasma glucose concentrations and glucose iAUC during an IV glucose tolerance test compared to 0.167 mg/kg/hr (n=3) and saline (n=6).

**Figure 3.**
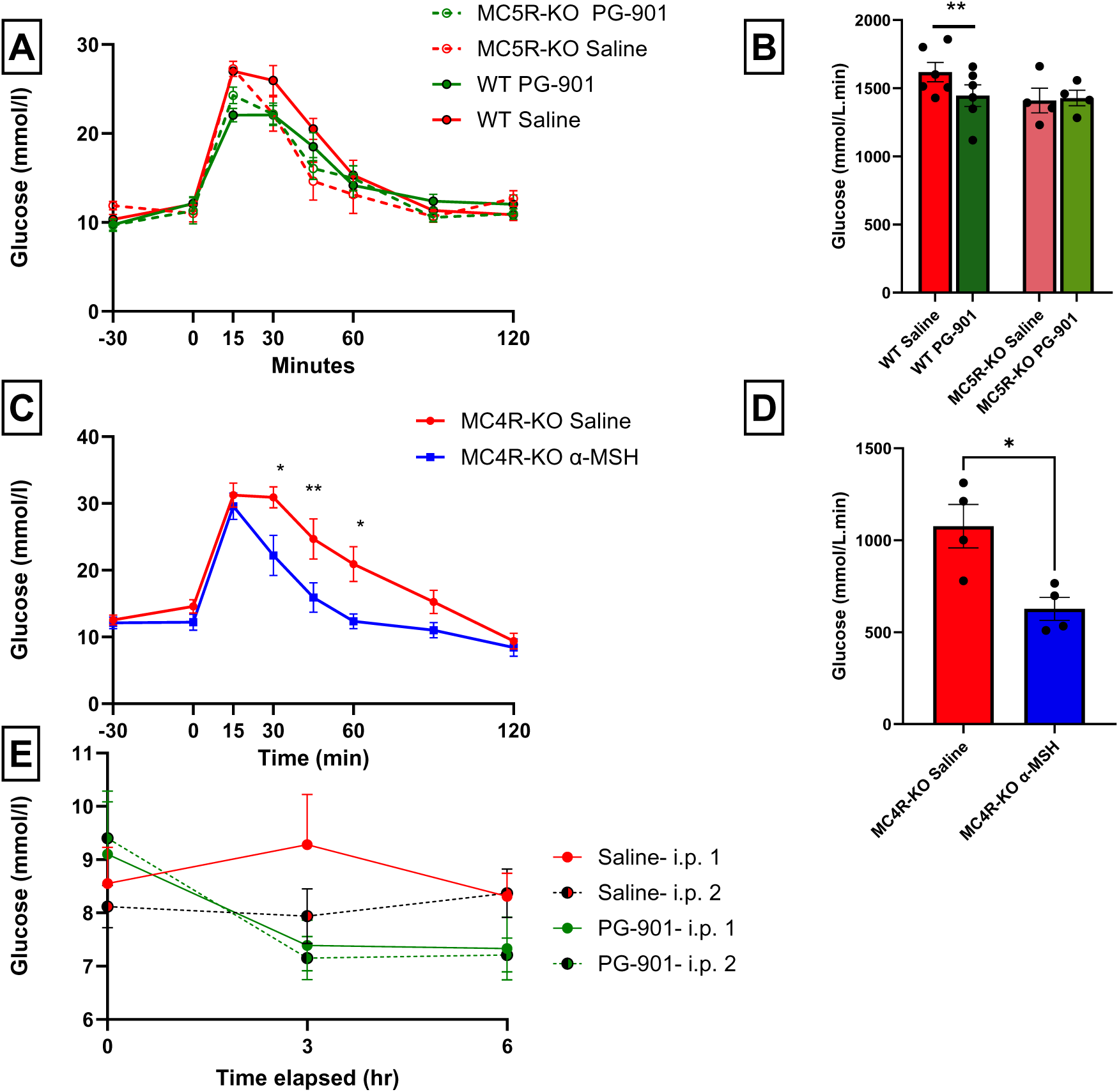
Effect of PG-901 and α-MSH in MC5R and MC4R KO mice. Data are presented as mean ± SEM.**A-B)** In male MC5R WT (n=6) or KO mice (n=4), ip. Injection of PG-901 or saline, followed by Glucose (2ug/g), in WT mice PG-901 reduced plasma glucose concentrations and glucose iAUC. **C-D)** In MC4R KO mice (n=4), mice IP treated with saline or αMSH (1ug/g), followed by glucose at 0mins, when MC4R KO mice were treated with αMSH, glucose tolerance was improved at 15, 30 and 60 mins post glucose and iAUC was significantly reduced. In MC4R KO mice, PG-901(1ug/g) treatment reduced blood glucose from baseline after both treatment 1 and treatment 2, no significant change was seen following saline (n=10).

Female southern pigtail macaques (*Macaca nemestrina*) were anaesthetized and treated via intravenous infusion with either saline or a selective MC5R agonist ^15^, PG-901 (0.167 or 0.334 mg/kg/hr), followed 30mins later with an intravenous glucose tolerance test (600mg/kg). In female pigtailed macaques, PG-901 demonstrated a non-significant increase in glucose tolerance, reducing glucose iAUC by 31% compared to saline at a dose of 0.334 mg/kg/hr (Figure 2 C-D).

### Role of MC5R in Glucose Tolerance in Mice

Glucose tolerance, as assessed by intraperitoneal glucose tolerance tests (IPGTTs), was significantly improved in wild type (WT) C57Bl6 mice, pre-treated with PG-901 (1µg/g of BW) 15 minutes prior to glucose injection, compared to vehicle-treated WT mice. At 15 minutes-post glucose bolus, plasma glucose concentrations were significantly lower in PG-901-treated animals compared to vehicle treated mice (22.0±0.8 mmol/l vs 27.0±0.3 mmol/l). iAUC for plasma glucose concentration in PG-901 treated mice was significantly lower than that of vehicle-treated mice (p<0.05). In IPGTTs performed in MC5R deficient mice^16^, there was no difference in glucose tolerance during treatment with PG-901 (1µg/g) compared to vehicle.

### Role of MC4R in Glucose Tolerance in Mice

The melanocortin 4 receptor plays an important role in weight homeostasis and has been implicated in glucose homeostasis so we tested the effect of peripheral administration of α-MSH and PG-901 in MC4R deficient mice^17^. α-MSH had no effect on plasma glucose when administered in the fasted state, but in IPGTTs in MC4R KO mice, pretreatment of α-MSH (1 µg/g) 30 minutes prior to glucose bolus administration significantly increased glucose tolerance compared to that of vehicle-treated mice.

In contrast with all other models studied in our experiments, in MC4R KO mice – which were mildly hyperglycemic, PG-901 reduced plasma glucose concentrations when administered in the fasted state, causing a 22% reduction in plasma glucose at 30 and 60 minutes. This effect was maintained over two consecutive injections of PG-901 (1 µg/g) administered six hours apart. There was no significant change in food intake or body weight during treatment with PG-901 compared to vehicle (**Supplementary Figure 4)**.

### Effect of α-MSH and PG-901 on glucose Uptake in Human Myotubes

Glucose uptake in immortalised human myotubes (differentiated from LHCN-M2 myoblasts^18^, a kind donation from Dr Mouly, Sorbonne Université, INSERM, Institut de Myologie, Paris, France) was quantified by high-performance anion-exchange chromatography with amperometric detection (HPAEC-PAD). The 2-Deoxy-D-glucose (2-DG) uptake was significantly increased by treatment with 100nM α-MSH for 1 hour compared to vehicle treated or untreated cells alone (270.7±79.65 nmol/mg vs. 312.2±63.71 nmol/mg) (**Fig 4. A**). When treated with the MC5R specific agonist PG-901 at 10nM and 100 nM, glucose uptake was also significantly increased in a dose dependent manner (270.7±79.65 nmol/mg vs. 344.2±115.6 nmol/mg and 393.8±111.9 nmol/mg, respectively) (**Fig 4. B)**.

**Figure 4.**
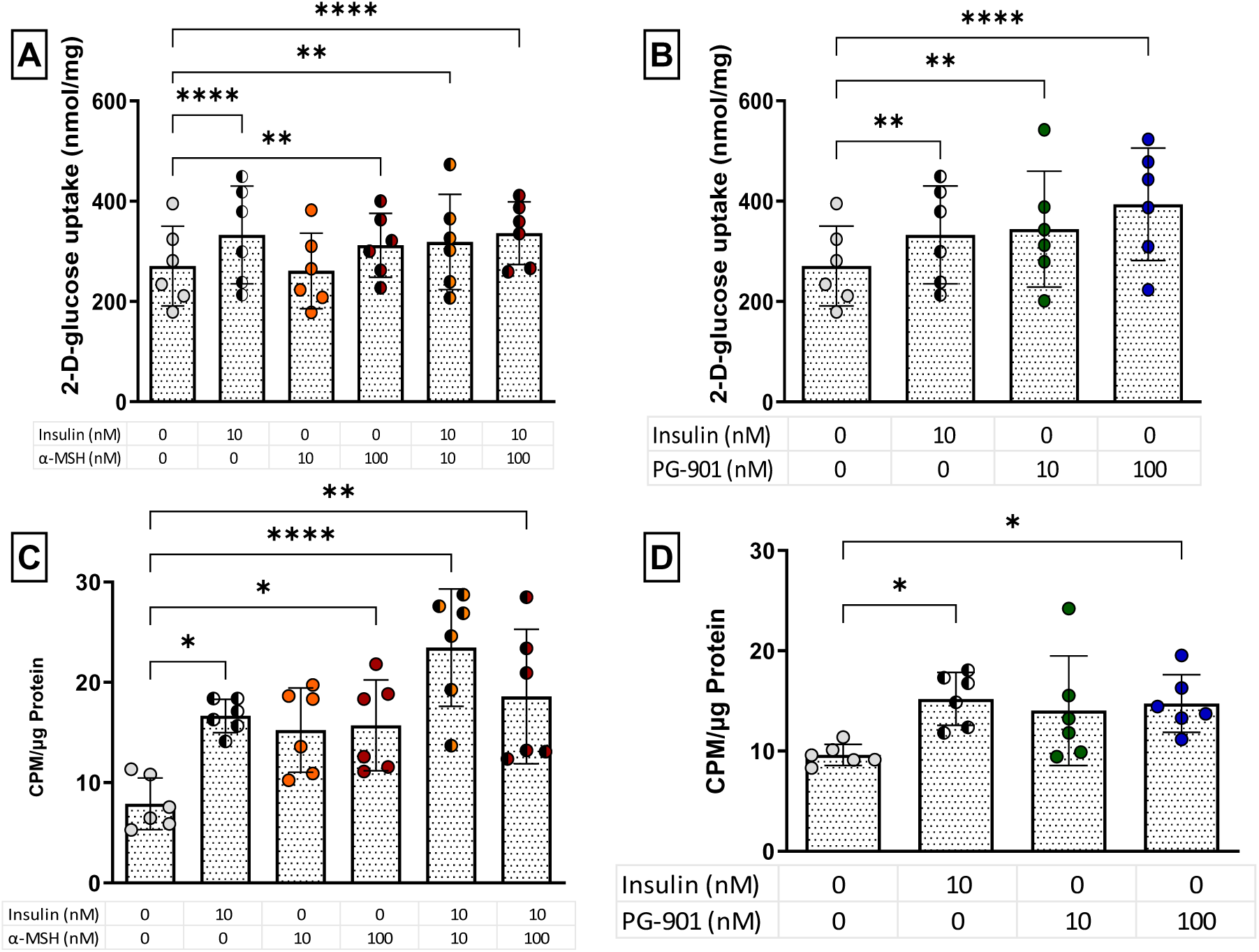
α-MSH and PG-901 administration induced glucose uptake in immortalised and primary human myotubes. **A)** Glucose uptake in immortalised LHCN-M2 myotubes in response to α-MSH and/or insulin. **B)** Glucose uptake in immortalised LHCN-M2 myotubes in response to the MC5R agonist PG-901 and/or insulin. **C)** Incubation with α-MSH induces glucose uptake in human primary myotubes. **D)** Incubation with the MC5R agonist PG-901 induces glucose uptake in human primary myotubes. Data are presented as mean ± SEM with individual data points from experiments performed in duplicate for each cell passage across three biological passages (n/N = 6/3). *p<0.05, **p<0.01, ***p<0.001, ***p<0.0001 assessed by repeated measures two-way ANOVA with post-hoc Tukey’s correction for multiple comparisons.

To test the validity of these results, primary human myoblast cell lines were established using *vastus lateralis* biopsies from healthy humans. Similar to data derived from the immortalised line, treatment of myotubes (differentiated myoblasts) with 100nM α-MSH significantly increased glucose uptake above untreated control (7.8±2.5 CPM/µg vs. 15.7±4.4 CPM/µg) (**Fig 4. C)**. The effect on glucose uptake was comparable in magnitude to that observed after treatment with the 10nM insulin (16.7±1.7 CPM/µg). When co-incubated with insulin, 10nM and 100nM α-MSH increases glucose uptake over insulin alone by 29.1% and 11% respectively. 100nM PG-901 also significantly increased glucose uptake over control in these cell lines (**Fig 4. D**).

## Discussion

This series of studies represent a multi-species and interdisciplinary approach to explore the role of the melanocortin system in glucose homeostasis. A series of human, non-human primate, rodent, and in vitro experiments demonstrated that α-MSH induces skeletal muscle glucose uptake manifesting physiologically as improved glucose tolerance. The mechanism underpinning this physiological observation involves skeletal muscle glucose uptake downstream of MC5R agonism and is independent and additive to insulin.

We have shown here in rodents, non-human primates, and humans that acute peripheral administration of α-MSH induces physiologically relevant reductions in plasma glucose. In humans, clamp experiments demonstrated that the mechanism underlying glucose lowering is peripheral uptake of plasma glucose. The simultaneous reduction in serum insulin is most likely a secondary phenomenon triggered by the sensing of reduced glucose concentrations by the pancreas. The lack of any difference in the incidence of hypoglycaemia between α-MSH and saline infusions is reassuring and suggests that insulin secretion remains appropriate for ambient glucose concentrations even at pharmacological concentrations of α-MSH in healthy humans. The lack of an effect of α-MSH in the fasted state (in multiple models), further supports its glucose-dependent mechanism of action.

In humans during and after infusion of α-MSH there was no signal of malaise, flushing, and nausea. None of the infusion studies were terminated early due to adverse events. Published data have suggested that agonism of central and peripheral MC3R and MC4R can modulate cardiovascular functions by increasing blood pressure and heart rate^19–21^, but our studies did not demonstrate any significant changes in these parameters. We assume this is because the peripherally administered α-MSH did not access CNS sites mediating hypertension. However, it is worth noting that bremelanotide, a non-specific melanocortin agonist prescribed to pre-menopausal women for generalized hypoactive sexual desire disorder, causes transient hypertension, and is contraindicated in patients with uncontrolled hypertension.

To determine whether selective MC5R agonism had a comparable effect to that observed by α-MSH, we extended our studies into non-human primates. Although constrained by small sample sizes, the results confirm that α-MSH and PG-901 increases glucose tolerance in non-human primates and strongly support the premise that MC5R is primarily responsible for the acute effect on glucose tolerance. In non-human primates, as with humans and normoglycemic mice, α-MSH and PG-901 had no effects on fasting glucose or insulin but reduced the glucose iAUC of glucose by 29% and 31% respectively. These data suggest that the effect can be achieved with MC5R agonism alone, without requiring parallel agonism at other melanocortin receptors. A peptidomimetic MC5R specific agonist, PG-901, did not alter glucose tolerance in MC5R deficient mice, but increased glucose tolerance in wild type mice and non-human primates. Previous studies have already demonstrated that α-MSH was not effective in changing glucose tolerance in MC5R deficient mice, despite increasing glucose tolerance in wild type mice^13^.

In MC4R deficient mice, α-MSH increased glucose tolerance, demonstrating that intact MC4R signaling was not required for the glucose lowering effect of the hormone. Despite not decreasing blood glucose in MC5R deficient mice, repeated doses of PG-901 lowered blood glucose in MC4R deficient mice following both injections, demonstrating that PG-901 is not acting as an antagonist at MC4R and that MC4R signaling was not required for increased glucose tolerance^15^. Additionally, a second dose of PG-901 continued to lower blood glucose demonstrating that the MC5R does not become rapidly desensitized or internalized^22^. Taken together, this set of in vivo pharmacological and molecular genetics studies show that the reduction in blood glucose caused by α-MSH is mediated through MC5R and is conserved across species.

Our series of *in vitro* experiments interrogated the cellular mechanism of action. In immortalised human skeletal muscle myotubes, 100nM α-MSH alone increased glucose uptake to a similar degree to 10nM insulin. When incubated with PG-901, a dose-dependent glucose uptake response was observed, showing that the MC5R-mediated glucose uptake response is conserved in human skeletal myotubes grown in culture. When incubated with α-MSH and PG-901, these cells displayed a similar glucose uptake response to the immortalised cells, confirming our findings in a primary cell model. These findings support our hypothesis that skeletal muscle is responsible for the post-prandial reduction of plasma glucose during α-MSH administration.

The skeletal muscle-mediated increase in glucose uptake appeared to be additive to the effect of insulin because co-administration of PG-901 and insulin caused a bigger increase in glucose uptake into immortalised muscle cells than either compound alone. This is consistent with the additive effect observed in ex vivo mouse soleus and L6 cells^13,14^. We have previously described that the mechanism by which α-MSH/MC5R affects glucose uptake does not involve Akt/Glut 4 and these new cell culture data supports this^14^. In our human studies, α-MSH infusion led to increased glucose tolerance, in spite of reduced insulin secretion. We propose that the combination of insulin-mediated and MC5R-mediated glucose uptake had an additive effect to that of insulin. This observation has clinical relevance, if agonism of the α-MSH axis has therapeutic potential as an insulin-sparing therapy in diabetes.

Across healthy humans, mice, and non-human primates, α-MSH and PG-901 had no effect on basal or fasting glucose concentrations. In contrast to all other results, in MC4R KO mice, PG-901, but not α-MSH, reduced glucose in the fasting state. This model had higher fasting blood glucose concentrations, so it is possible that MC5R-induced glucose tolerance acts only on higher blood glucose levels. We have previously proposed that MC5R glucose tolerance is mediated through a low-affinity glucose transporter^14^, which could be a plausible explanation.

While all models used are in concordance with respect to skeletal muscle glucose uptake and whole-body glucose tolerance, we acknowledge the series of studies presented have some limitations. As outlined previously, MC5R biology in humans is less well explored relative to the other melanocortin receptors. There are no validated antibodies for use in muscle cells, no available ligands that provide receptor labelling, and a lack of highly specific antagonists to demonstrate disruption of receptor functionality. In rodents, hypothalamic MC4R has been linked to increased glucose disposal and reduced glucose production^11^, as well as glucose uptake into intrascapular brown adipose tissue^12^. Therefore, the observation that α-MSH improves glucose tolerance in healthy humans, could also be mediated in parallel by activation of hypothalamic MC4R. However, in mice, α-MSH still lowers glucose in mice without MC4Rs. The results in non-human primates suggest that MC5R may be the primary contributor to improve glucose tolerance, however these data were obtained from a small number of non-human primates. Despite these limitations, the multi-species and multi-site nature of these series of studies provide compelling evidence for the key role of the α-MSH/MC5R pathway in glucose metabolism. Future studies are planned to fully elucidate the intracellular signaling pathway downstream of MC5R.

Through this series of translational studies, we have shown that a) infusions of α-MSH in healthy humans increased glucose tolerance, b) treatment of human muscle cells *in vitro* with the peptide induced a glucose uptake response, and c) this effect is replicated by a selective MC5R agonist inducing glucose uptake into human muscle cells in vitro and improving glucose tolerance in non-human primates and mice. This novel physiological response to melanocortin receptor agonism in humans, implicates a melanocortin-skeletal muscle centric pathway which mirrors our previously published pre-clinical data. Manipulation of this pathway may have therapeutic potential for metabolic diseases such as diabetes mellitus.

## Methods

### Sex as a biological variable

For human studies male and female participants were recruited as equally however due to the odd N number of the dose-finding study cohort the number of male participants is one greater than the female participants. In the animal studies, male mice and both male and female macaques were included. However, due to limitations in the number of macaques and sexes available during the study period, a complete comparison of all drugs and doses across both sexes could not be conducted. While this may be limiting, we have made every effort to ensure representation of both sexes.

### Human studies

#### Participants

Participants for the human infusion studies were recruited from the community and Imperial Clinical Research Facility Healthy Volunteer Database. Participants were eligible if they were male or female, aged 18-50 years, and had a BMI ≥18 and < 30 kg/m^2^. Key exclusion criteria were the presence of any form of diabetes or other significant physical or mental health disease (Full study protocol included as supplementary file).

#### Infusion Study Design

A preliminary, dose-finding study with 15 participants was conducted to determine (i) whether acute administration of α-MSH lowers blood glucose in a dose-dependent manner, (ii) the effect size of the α-MSH dose with the greatest impact on glucose-lowering and (iii) the safety profile of acute intravenous α-MSH infusions. Following screening, participants were invited to attend the NIHR Imperial Clinical Research Facility 6 times. The first 4 visits comprised of OGTTs carried out during infusions of saline or one of 3 doses of α-MSH; low (15 ng/kg/hr), medium (150 ng/kg/hr) and high (1500 ng/kg/hr) in a placebo controlled, randomized, double-blinded manner. The following two visits comprised of hyperinsulinaemic-euglycaemic clamps carried out with an insulin infusion of 1 µU/kg/hr with a concomitant infusion of saline or 150 ng/kg/hr α-MSH. These visits were also double-blinded and randomised.

The effect size measured in the dose-finding OGTT study was used to estimate the sample size of the replication study in which the effect of acute administration of α-MSH vs saline was compared The α-MSH used in these studies was synthesized by Auspep Clinical Peptides Ltd (Australia) to GMP standards. PG-901 was synthesized using Fmoc-based solid phase synthesis methods for the linear sequence followed by solution cyclisation using PyClock reagent^15^. The product was purified by RP-HPLC.

#### Experimental Procedures

##### Dose-finding study

Fifteen healthy participants were asked to refrain from alcohol and strenuous physical activity for 48 hours before the study visits. They were also asked to consume a standardised meal the evening before the study and only consume water from 10pm onwards. Female participants were studied in the follicular stage of their menstrual cycle.

##### Replication study

A different cohort of 22 healthy human participants were recruited to this study from the community and a healthy volunteer database. The same methods used in the dose-finding study were followed. Participants attended on two occasions where they underwent a standard OGTT whilst receiving the high dose (1500ng/kg/hr) of α-MSH or saline in a randomised double-blinded manner.

##### Oral glucose tolerance tests

Upon arrival, participants were cannulated in the antecubital vein of each arm then allowed to rest for 30 minutes, after which they received an infusion of either saline or low (15ng/kg/hr), medium (150mg/kg/hr), or high doses (1500mg/kg/hr) of α-MSH. The order of the infusions was random and both investigators and participants were blinded to the contents of the infusion. Thirty minutes after the infusion had started patients underwent a standard OGTT by consuming 75g of dextrose dissolved in 300ml of water. Blood samples were collected at the following time points relative to initiation of infusion: −30, 0, 15, 30, 60, 120 minutes. Participants were also asked to complete appetite and safety visual analogue scales at the same time points.

##### Euglycaemic-hyperinsulinaemic clamps

For the remaining two visits, participants received an infusion of either saline or medium dose of α-MSH during a euglycaemic-hyperinsulinaemic clamp. The order of the infusions was random and both investigators and participants blinded to the contents of the infusion. During the clamp, insulin was infused at a rate of 1mU/kg/min for 180 minutes. Euglycaemia was maintained by infusing 20% dextrose at a variable rate. The glucose infusion rate (GIR) at the plateau of the clamp (minutes 150-180) was used as a marker of glucose uptake from plasma.

##### Sample Analyses

Plasma samples collected at the OGTTs and clamps were separated immediately by centrifugation at 4°C and then stored at −80°C until analysis. Serum samples were allowed to clot for 30 minutes before centrifugation. Plasma glucose analysis was conducted at the Imperial College Healthcare NHS Trust pathology laboratory using the hexokinase method. Serum insulin was measured using the Alinity reagent kit (4T7520) on an Abbott Architect i2000SR. Plasma α-MSH concentration was measured using radioimmunoassay (coefficient of variation 12.5%, Phoenix Pharmaceuticals; CA, USA).

##### Statistical considerations

The aim of the dose-finding study was to determine the effect size of the α-MSH dose with the highest glucose-lowering effect. Thus, due to the exploratory nature of the study, no formal statistical comparisons were performed.

The high dose of α-MSH had the greatest glucose-lowering effect (**Supplementary Table 2**) and was not associated with any adverse events. Based on that, it was estimated that 22 participants for the Replication study were required to provide 80% power to detect a statistically significant difference between α-MSH at a dose of 1500 ng/kg/hr vs. saline at α=0.05.

Data were tested for normality distribution and summarised using descriptive statistics. Differences in metabolic outcomes between saline and the high dose of α-MSH were compared using paired t-tests and repeated measures 2-way ANOVA using the Šídák correction for multiple comparisons.

Data are reported as mean ± SEM unless stated otherwise. iAUC was calculated with the trapezoidal rule, correcting for baseline with the glucose measurement at timepoint 0. All analysis were performed with GraphPad Prism 8.0.

##### Ethical Approvals

Human infusion studies were conducted at NIHR Imperial Clinical Research Facility (London) and approved by the London-Fulham Research Ethics committee (20/LO/0355, ISRCTN 26265936). Muscle biopsy studies were performed in University College Dublin and approved by the UCD Human Research Ethics Committee (LS-18-101-Docherty). All participants provided written informed consent prior to participation. These studies were conducted in line with the Declaration of Helsinki (2008).

## Animal studies

### Non-Human Primate Studies

#### Southern Pigtailed Macaques and Ethical Approval

Southern pigtail macaques (*Macaca nemestrina*) used in the study were procured from the National Non-Human Primate Research and Breeding Facility (Monash University). Animals were housed in single or double Flex-A-Gon cages (Britz and Company, USA) in single-sex groups containing 5-10 animals per enclosure with indoor and outdoor access. Animals were fed standard primate chow (Gordons Specialty Feeds, Australia) once daily (200g/macaque) with an energy density of 13Mj/kg (21% protein, 5.3 % fibre, and 8% fat). Animals also received daily seasonal fruits and vegetables (300g/animal) in addition to food items provided as part of the facilities enrichment program (dried fruits, nuts, seeds, maize, jelly, fruit juice, treacle, muffins, popcorn, meal worms, coconut flakes and plant foliage). All animals had ad libitum access to water. A total of 9 male and 11 female Southern pigtail macaques, aged 4-17 years were studied.

#### Infusion Study Design - Intravenous Glucose Tolerance Test (ivGTT)

Animals fasted overnight were transferred from group housing into modular primate housing units (PHUs) and restrained with the aid of the squeeze back system. Animals then received a single intramuscular injection (gluteal or quadricep muscle) of an equal mixture of tiletamine/zolazepam (1.4–3.2 mg/kg Zoletil100; Virbac Animal Health Australia) combined with butorphanol (0.1– 0.3mg/kg; Butorphanol Tartrate (Butrogesic), Troy Laboratories Australia) and atropine (0.02– 0.04mg/kg; Atropine Sulphate (Atrosite (Illium)), Troy Laboratories Australia). Once immobilised, the animal was then transported to the procedure room, placed onto a pre-heated, fan-forced hot-air filled blanket (Darvall, Advanced Anaesthesia Specialists) and maintained on 2-4% Sevoflurane in 100% oxygen via a face musk until diminished muscle tone, loss of palpebral reflexes, pedal reflexes were noted and pulse and respiration rates were stabilised. The animal was then intubated with a cuffed endotracheal tube and ventilated for the entire duration of the procedures with 1-2% Sevoflurane in 100% oxygen in a prone position. An indwelling iv catheter-18G, 1.3 x 48mm (BD, USA) was inserted into the superficial saphenous vein (caudal region) and used to infuse saline (vehicle), α-MSH (0.167 or 0.334mg/kg/hr (Auspep, Australia)) or PG-901 (0.167 or 0.334mg/kg/hr) for 30 minutes prior to the bolus administration of 50% glucose solution (600mg/kg). Blood sampling at −30, −15, 0, 1, 3, 5, 10, 20, 40 and 60 minutes was performed using a 27G X ½” needle on the adjacent limb. Saline (vehicle), α-MSH (0.1667-0.334mg/kg/hr) or PG-901 (0.1667-0.334mg/kg/hr) infusion was maintained throughout the entire duration of the ivGTT. Glucose in whole blood was immediately measured using an Accu-Chek Performa blood glucose monitor (Roche Diagnostics).

#### Mouse experiments

Lean 8 week old male WT or MC5R deficient mice^13^ were treated intraperitoneally (ip) with either vehicle (veh) or PG-901 (1 μg/g of body weight). Fifteen minutes after the injection, the mice received a bolus of glucose (2 μg/g of body weight). Blood samples were collected from a tail nick to measure glucose concentrations.

Male MC4R KO (loxTB Mc4r, C57BL/6J, Jax: 032518) mice were injected IP with α-MSH (1 μg/g) or vehicle. Thirty minutes later, the mice were injected IP with glucose (1 μg/g of body weight). Blood samples were collected to measure glucose concentrations.

Male MC4R KO mice were treated IP with PG-901(1 μg/g) or vehicle. Blood glucose concentrations were measured at 0, 30, and 60 minutes. To assess if there was any tachyphylaxis to MC5R agonism six hours after the first injection of vehicle or PG-901, the mice were reinjected with PG-901, and blood glucose concentrations, food intake and body weight were measured at the same time points.

#### Ethical Approval for *In Vivo* Studies

All animal protocols, health and wellbeing monitoring guidelines including experimental endpoints were approved by the Monash University Animal Ethics Committee (AEC) in accordance with the Australian Code for the Care and Use of Animals for Scientific Purposes, 8th Edition, 2013 and the National Health and Medical Research Council Principles and Guidelines for the Care and Use of Non-Human Primates for Scientific Purposes, 2016.

### In vitro studies

#### Primary Myoblast Isolation

Muscle biopsies taken from the *vastus lateralis* of healthy volunteers were used to establish primary myoblast cultures according to published protocol^23^. Upon arrival, participants were allowed to rest and administered a subcutaneous 2% lidocaine solution as a local anaesthetic. A 14G cannula was inserted at an angle directly into the *vastus lateralis*. A co-axial spring-loaded needle was then inserted into the cannula to obtained muscle biopsies of approximately 50mg in weight. The muscle biopsies were immediately placed in DPBS (Merck, Darmstadt, DE) on ice and shortly after minced with a scalpel and dissociated with enzymatic solution (DPBS containing 10 mg/mL Collagenase D, 4.8 mg/mL Dispase II, and 2.5 mM CaCl_2_)…at 37oC for 40 minutes.

The solution was then transferred to a 50mL falcon tube and incubated at 37°C for 40 minutes with gentle agitation. Skeletal Muscle Growth Medium (Cell Applications Inc., San Diego, USA) was then added to the enzymatic solution which was then passed through a 70 µM cell strainer and centrifuged at 443g for 5 minutes at room temperature. The isolated cell pellet resuspended in Skeletal Muscle Growth medium and plated in collagen coated plates.

#### *In vitro* Glucose Uptake

LHCN-M2 immortalised human skeletal myoblasts were cultured and differentiated to form myotubes (72 h) as we have done previously^24^. Myotubes were starved of serum and glucose for 4 h, then incubated with insulin (Actrapid, Novo Nordisk) and/or α-MSH or PG-901, or vehicle (untreated control), and 2-deoxy-D-gluose (2-DG) for 30 min. The incubation media were collected and snap-frozen. Cells were washed, lysed and total protein measured by BCA. Media were prepared for and analysed by HPAEC-PAD, following our established protocol^25^. HPAEC-PAD peak areas for 2-DG in the media were compared to the standard curve (**Supplementary Figure 5**), subtracted from the initial 2-DG concentration and corrected for total protein of the corresponding cell lysate. Data are presented as intracellular uptake of 2-DG. Since data were all normally distributed and heteroscedastic, and residuals were also normally distributed, two-way ANOVA (with insulin and α-MSH or PG-901 as variables matched across each experiment) was used to assess significant differences between the treatment groups with post-hoc Tukey’s multiple comparisons.

Primary human myoblasts derived from dispersed vastus lateralis muscle biopsies from healthy humans were cultured and differentiated to form multinucleated mature myotubes in 24-well plates. To differentiate, primary myoblasts were grown to confluency then incubated with Skeletal Muscle Differentiation Media (PromoCell, Heidelberg, DE), changing media every 2 days. After 5-7 days when multinucleated differentiated myotubes were observed, cells were assayed for glucose uptake. Prior to the assay, myotubes were serum starved in low-glucose (1 g/L) DMEM for 6 hours before being glucose starved for 1 hour in glucose-free DMEM. Myotubes were then incubated with glucose and serum free DMEM (control), 10nm recombinant human insulin (Roche Life Science, Basel, CH), or PG-901 for 1 hour. Tritiated 2-deoxy-D-glucose [3H], 2-deoxy-D-glucose (PerkinElmer, Waltham, USA) was mixed with unlabeled 2-deoxy-D-glucose then added to directly to the treatment solution in each well for a final concentration of 5.5 mM glucose and 0.5 µCi/well, then incubated for 10 minutes. Cells were placed on ice, washed with ice-cold PBS, and lysed with Cell Lytic MT™ (Merck. Darmstadt, DE) for 15 minutes. Lysate was then mixed with Ultima Gold™ scintillation cocktail and counts per minute (1 minute/sample) were measured in a liquid scintillation counter. An aliquot of lysate was reserved for protein quantification and glucose uptake expressed as counts per minute per microgram of protein counted.

## Data Availability

Please contact the corresponding authors.

## Author Contributions

PS, BJ and SES contributed equally to this work. This work comprises equally important in vitro, human and in vivo aspects driven respectively by the first authors. The order the co-first authors are listed was determined by a random sequence generator.

PS acquired, analysed, and interpreted the *in vitro* myocyte data, contributed substantially to the acquisition, analysis and interpretation of the dose-finding study data, and preparation of the manuscript.

BJ contributed substantially to the acquisition, analysis and interpretation of the dose-finding and replication study data, and preparation of the manuscript.

SES contributed substantially to the acquisition, analysis, and interpretation of *in vivo* data.

TM contributed substantially to the acquisition, analysis, and interpretation of *in vivo* data.

MJH conducted glucose uptake studies in LHCN-M2 immortalised human skeletal myotubes.

SS contributed substantially to the acquisition, analysis and interpretation of the dose-finding and replication study data.

NGD contributed substantially to study design and data interpretation and manuscript preparation.

JTP contributed to analysis of in vivo animal data.

CG contributed to the conduct of the study and management across sites.

AG conducted in vivo animal studies.

PET synthesized PG 901

WKY synthesized PG 901

GdV collected the muscle biopsies from human subjects.

GW contributed to glucose uptake studies in LHCN-M2 immortalised human skeletal myotubes.

CWlR made substantial contributions to the conception and design of the study, critically reviewed the manuscript, and provided final approval of the version to be published.

ADM made substantial contributions to the conception and design of the study, critically reviewed the manuscript, and provided final approval of the version to be published.

MAC made substantial contributions to the conception and design of the study, critically reviewed the manuscript, and provided final approval of the version to be published.

## Acknowledgements

This work was supported by; the Health Research Board Ireland (ILP-HSR-2017-007), European Foundation for the Study of Diabetes (Travel Fellowship), National Health and Medical Research Council (Australia) Victorian Medical Research Accelerator Fund, and Breakthrough T1D (formerly JDRF International) (Strategic Research Agreement). The funders had no role in study design, data collection and analysis, decision to publish, or preparation of the paper.

We thank Monash Animal Research Platform for overseeing the husbandry and care of the non-human primates and rodents used in the studies and for providing technical assistance in performing the described glucose tolerance tests performed in non-human primates.

This paper presents independent research funded by grants from the National Institute for Health and Care Research (NIHR) and supported by the NIHR Imperial Clinical Research Facility and NIHR Imperial Biomedical Research Centre. The Section of Endocrinology and Investigative Medicine is funded by grants from the Medical Research Council (MRC) and the NIHR. The views expressed are those of the authors and not necessarily those of the MRC, the NHS, the NIHR, or the Department of Health.

