## Supplemental Data and Figures for "α-Melanocyte Stimulating Hormone Reduces Blood Glucose Across Species"

### Supplemental Figures

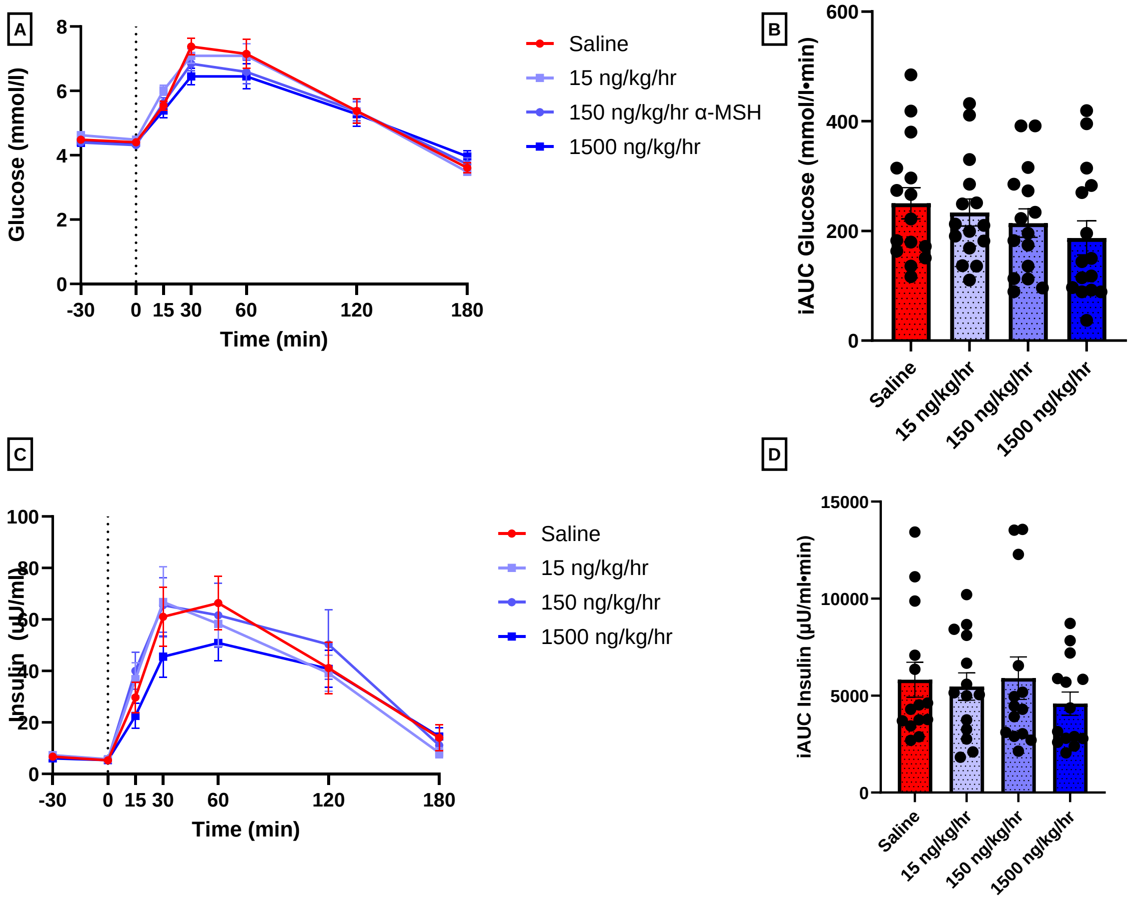


**Supplemental Figure 1.** Effect of intravenous infusion of α-MSH (15, 150, and 1500 ng/kg/hr) in healthy humans in the dose finding cohort. Data are presented as mean ± SEM. Timepoint of oral glucose administration marked by dotted line. A-B) A dose of 1500 ng/kg/hr α-MSH caused the greatest reduction in plasma glucose during an OGTT (n=15). B-C) A dose of 1500 ng/kg/hr α-MSH caused the greatest reduction in serum insulin during an OGTT(n=15).

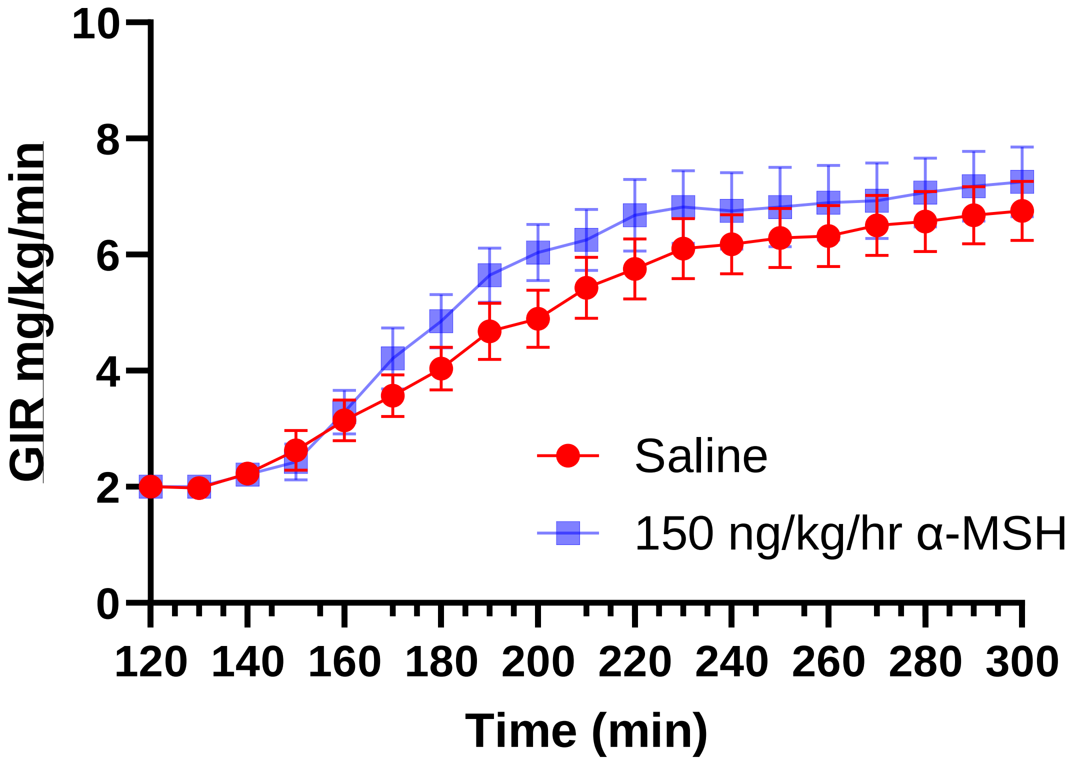


**Supplemental Figure 2.** Intravenous infusion of 150 ng/kg/hr α-MSH during a hyperinsulinaemic-euglycaemic clamp in healthy humans (n=14). Data are presented as mean ± SEM

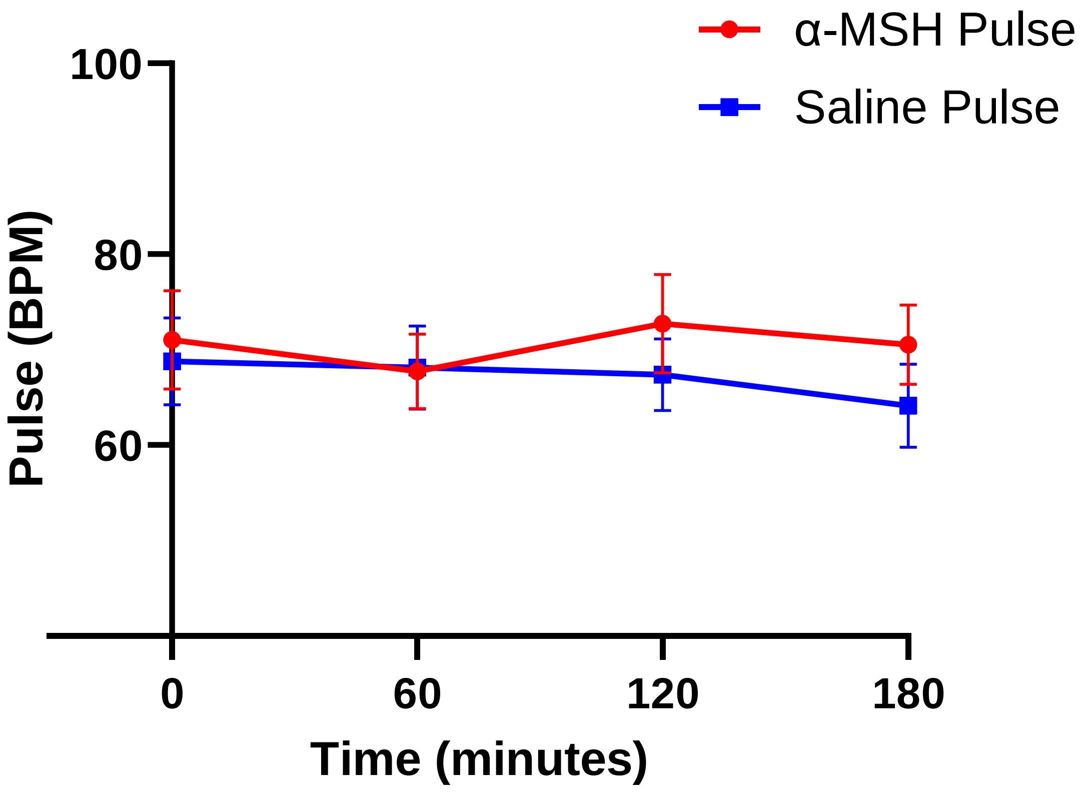


**<u>Supplemental Figure 3.</u>** Pulse (beats per minute) as measured by blood pressure monitor in healthy humans during intravenous α-MSH infusion (n=15). Data are presented as mean ± SEM.

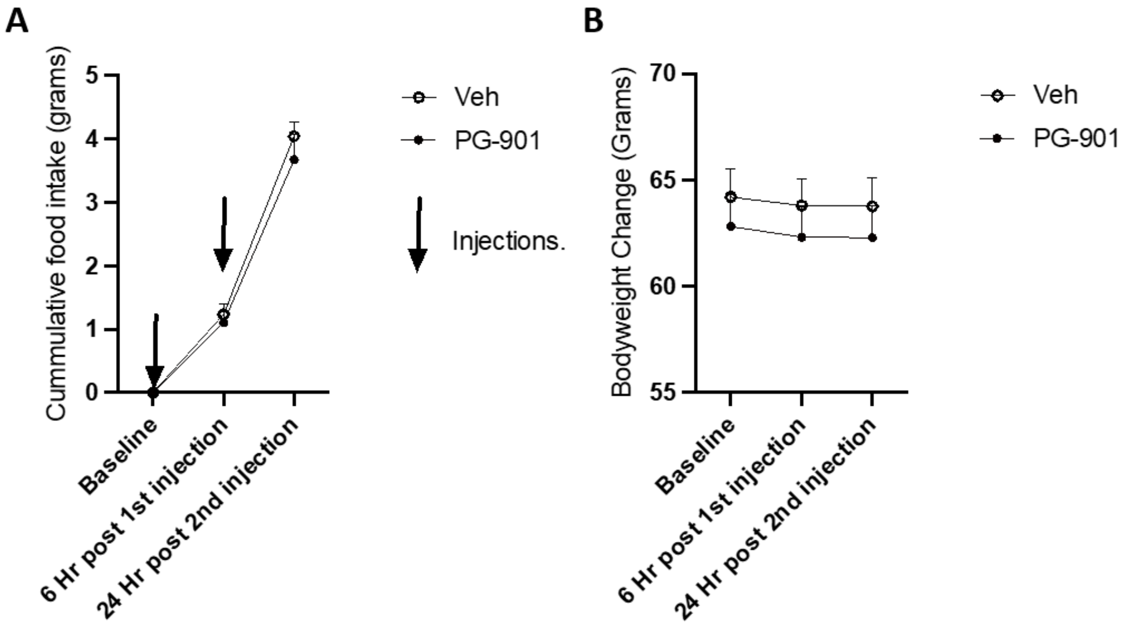


**Supplemental Figure 4.** Effect of PG-901 treatment in MC4R KO mice. Data are presented as mean ± SEM.**A-B)** In male MC4R KO mice, PG-901(1ug/g) treatment did not change cumulative food intake compared to vehicle treatment (A). In male MC4R KO mice, PG-901(1ug/g) treatment did not affect body weight compared to vehicle treatment (A) (n=10). Mean +/- SEM, n=10, Two way ANOVA, Tukey's multiple comparisons test.

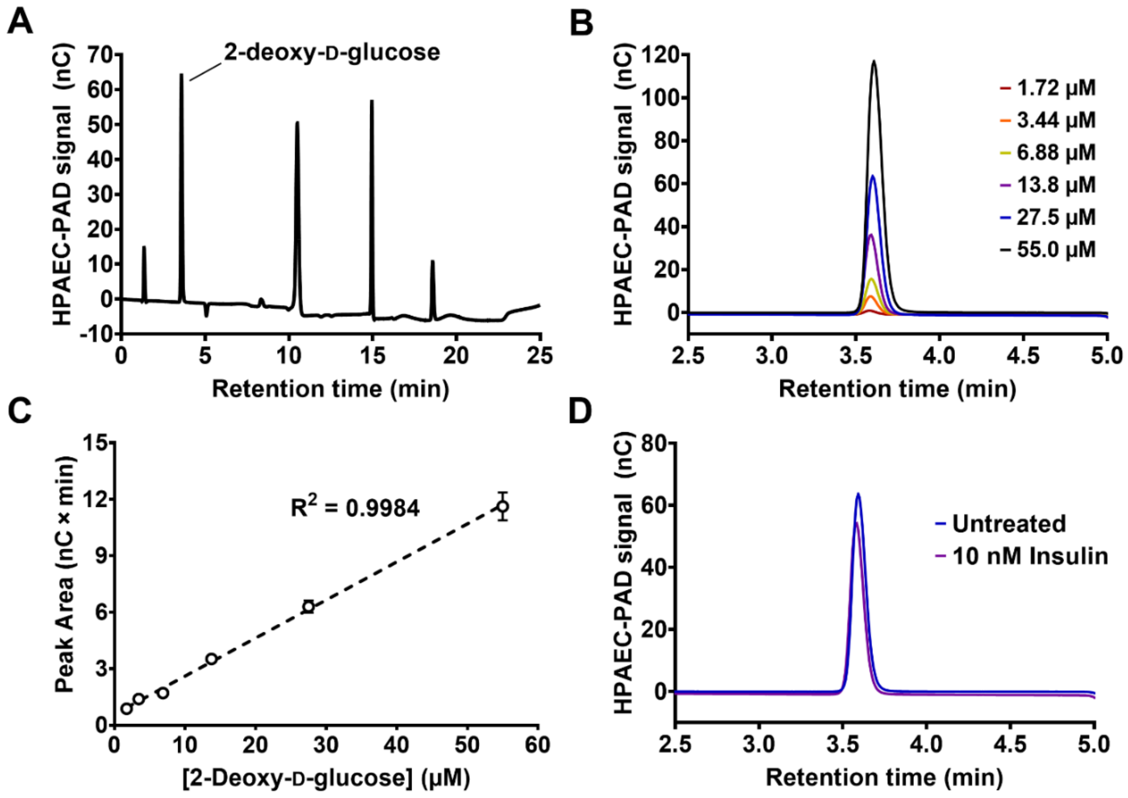


**<u>Supplementary Figure 5.</u>** Glucose uptake in immortalised human LHCN-M2 myotubes was measured by quantifying the intracellular uptake of 2-deoxy-D-glucose by high-performance anion-exchange chromatography with pulsed amperometric detection (HPAEC-PAD). **A)** HPAEC-PAD chromatogram for DMEM cell culture medium supplemented with 0.55 mM 2-deoxy-D-glucose. **B)** HPAEC-PAD chromatogram (only 2.5 - 5 min retention time shown), demonstrating peaks for varying concentrations of 2-deoxy-D-gluose reference standard in DMEM cell culture medium. **C)** Standard curve for 2-deoxy-D-gluose reference standards shown in B) (n = 3 injected in duplicate, mean ± SD). **D)** HPAEC-PAD chromatograms (only 2.5 - 5 min retention time shown) demonstrating peaks for example injections of media from LHCN-M2 myotubes after incubation ± insulin (10 nM) and 0.55 mM 2-deoxy-D-glucose for 30 min.

**Supplementary Tables**

| | Dose finding (n=15) | Replication (n=22) |
|---|---|---|
| **Sex** | **8M, 7F** | **11M, 11F** |
| **Age (years)** | **28 ± 8.4** | **29.1 ± 9.1** |
| **Weight (kg)** | **72.9 ± 14.6** | **70.3 ± 13.2** |
| **Body fat (%)** | **24.3 ± 5.9** | **21.4 ± 7.7** |
| **BMI (kg/m$^2$)** | **23.9 ± 3.0** | **22.7 ± 3.1** |
| **Fasting glucose (mmol/l)** | **4.5 ± 0.4** | **4.5 ± 0.4** |
| **HOMA-IR** | **1.4 ± 0.6** | **1.0 ± 0.5** |
| **HbA1c (mmol/mol)** | **32 ± 4.1** | **33 ± 0.8** |

**Supplementary Table 1**. Baseline characteristics for Dose Finding and Replication cohorts.

Data are presented as mean ± SD.

| Dose-finding Cohort (n=15)<br><br>Oral Glucose Tolerance tests | | | | | | | |
|---|---|---|---|---|---|---|---|
| | Incremental Area Under the Curve 0-120 minutes | | | | Mean difference vs. Saline<br><br>(95% Confidence Interval) | | |
| | Saline | 15 ng/kg/hr α-MSH | 150 ng/kg/hr α-MSH | 1500 ng/kg/hr α-MSH | 15 ng/kg/hr α-MSH | 150 ng/kg/hr α-MSH | 1500 ng/kg/hr α-MSH |
| Plasma glucose (mmol/l.min) | 243.9 ± 119.7 | 229.3 ± 99.6 | 212.5 ± 102.0 | 184.0 ± 123.4 | 14.6<br><br>(-47.5 to 76.6) | 31.4<br><br>(-31.4 to 94.2) | 59.9<br><br>(-2.1 to 121.8) |
| Serum insulin (µU/ml.min) | 5611 ± 3358 | 5004 ± 2713 | 5792 ± 3889 | 4577 ± 2171 | 606.7<br><br>(-380.2 to -1593.6) | -181.5<br><br>(836.0 - 472.9) | 1033.7<br><br>(-111.0 to 2178.4) |

**Supplementary Table 2.** Effect of intravenous infusion of α-MSH (15, 150, and 1500 ng/kg/hr) on plasma glucose and serum insulin in healthy humans (dose finding cohort). Data are presented as mean ± SD and mean difference vs. saline (95% confidence interval). N=15.

| Dose-finding Cohort (n=15) | | | | | | | |
|---|---|---|---|---|---|---|---|
| Visual Analogue Scale rating | Infusion | | | | Mean difference vs. Saline (95% Confidence Interval) | | |
| | Saline | 15 ng/kg/hr α-MSH | 150 ng/kg/hr α-MSH | 1500 ng/kg/hr α-MSH | 15 ng/kg/hr α-MSH | 150 ng/kg/hr α-MSH | 1500 ng/kg/hr α-MSH |
| Hunger (mm) | 44.9 ± 24.4 | 39.2 ± 20.8 | 36.7 ± 22.4 | 35.2 ± 19.0 | 5.7 (-7.2 to 18.6) | 8.2 (-5.1 to 21.4) | 9.7 (-7.1 to 26.5) |
| Pleasantness to eat (mm) | 53.2 ± 22.1 | 50.7 ± 25.2 | 45.8 ± 21.5 | 46.3 ± 22.6 | 2.5 (-12.4 to 17.4) | 7.4 (-3.9 to 18.8) | 6.9 (-8.0 to 21.9) |
| Fullness (mm) | 22.1 ± 16.9 | 25.6 ± 24.0 | 27.1 ± 22.25 | 28.4 ± 22.3 | 2.5 (-8.6 to 15.6) | 5.1 (-7.4 to 17.5) | 6.4 (-14.5 to 27.2) |

**Supplementary Table 3.** Sensations of Hunger, Pleasantness to eat, and Fullness as measured by Visual Analogue Scale (VAS) during intravenous α-MSH infusions in healthy volunteers (n=15). Data are presented as mean ± SD.
